# Proliferative events ameliorate DNA damage accumulation without affecting function in hematopoietic stem cells

**DOI:** 10.1101/2024.08.21.608631

**Authors:** Shubham Haribhau Mehatre, Harsh Agrawal, Irene Mariam Roy, Sarah Schouteden, Satish Khurana

**Author notes:** Corresponding author **Correspondence:** Satish Khurana, PhD, Associate Professor, School of Biology, Indian Institute of Science Education and Research Thiruvananthapuram, Kerala, India 695551.

## Abstract

Upon aging, HSCs show functional decline with increased proliferation, myeloid skewing, and poor engraftment efficiency. Accumulation of DNA damage has been causally linked with this phenomenon, with the debatable role of proliferative events. In this study, we sought to enquire the effect of increased hematopoietic stem cell (HSC) proliferation during the lifetime on the hematopoietic aging in mice. Multiple rounds of blood withdrawals were performed between two to twelve months of adult life to maintain higher proliferation rate in HSC population. Our experiments showed little effect of increased proliferation rate on age-associated functional decline in hematopoietic system. However, we noted a decrease in the double strand breaks (DSBs) accumulated with age in mice that underwent serial bleeding regimen. Analysis of single-cell sequencing data from mouse and human HSPCs showed enrichment of DNA damage response pathways confirmed by increased expression of the genes involved. Importantly, we demonstrate that the induction of HSC proliferation in aged mice is sufficient to decrease the load of DSBs. Hence, our results show that proliferative events during lifetime might aid in clearing age-associated DSBs. While these DNA damages might not be directly linked with the functional decline, proliferation induced clearance can have clinical implications.

## Main text

The number of long-term (LT-) repopulating HSCs with multi-lineage engraftment potential decreases drastically in aged mice and humans (*1*). Increased proliferation rates, such as in the case of deletion of cell cycle regulators p21, p27, and Bmi1, resulted in pre-mature aging like phenotype (*2*). DNA damage accumulation has been implicated with age-associated and proliferation induced functional decline in HSCs (*3*). Mice deficient in *Atm* expression (*Atm*^*−/−*^), that mediates response to DNA double strand breaks (DSBs), showed progressive bone marrow (BM) failure due to poorly functioning HSCs (*4*). The exit from quiescence in HSCs was directly linked to DNA damage and functional decline (*5, 6*). Contrarily, aged HSCs exhibited activation of DNA damage response (DDR) mechanisms and clearance of DNA damage upon cell cycle entry (*7*). Here, we sought to examine the effect of repeated proliferative events during the lifetime on age-associated functional decline in HSCs and DNA damage accumulation.

As blood loss is a known inducer of HSC proliferation (*8*), we used a serial bleeding-based regimen to test the effect of physiological demand driven proliferation on HSC aging (Schematic in Fig. 1a). We compared the blood cell counts in aged mice that underwent serial bleeding (old donors) with the age-matched (old controls; 12, 18 and 24 months) and young control mice. At 12 months, we did not observe any difference in WBC, lymphocyte, granulocyte, hematocrit, monocytes, eosinophil, RBC and hemoglobin levels between old control and old donor groups (Fig. S1a-h). However, a decrease in WBC, granulocyte and monocyte counts was noticeable in the two groups of old mice when compared with young controls (Fig. S1a,c,e). Even after 18 months of age, we did not notice any change in blood cell counts from old donors when compared with old control (Fig. S1i-p). Again, when compared to the young control mice, both of the old mice groups showed significant decline in WBC, lymphocyte, hematocrit, monocytes and hemoglobin levels (Fig. S1i,j,l,m,p) and a significant increase in granulocyte and eosinophils (Fig. S1k,n). At 24 months of age, an unchanged number of monocytes (Fig. S1q) and granulocytes (Fig. S1r) was observed with a decreased number of WBCs (Fig. 1b). Decrease in the WBC counts was reflected in lymphocyte counts (Fig. 1c) with a more robust change in old donor mice than old controls. We could also note a consistent decline in erythropoietic activity as there was a decrease in RBC numbers (Fig. 1d), along with hematocrit (Fig. 1e) and hemoglobin levels (Fig. S1s). Higher eosinophil numbers (Fig. 1f) could be linked with pro-inflammatory state that is known to get established with aging in mice and humans (*9*). With age, decreased erythrocyte output has been reported (*10*) along with an increase in the erythro-myeloid precursor cells (*11*). However, only 5-7% of individuals aged >65 years were found mildly anemic (*12*). This age-associated anemic state could also be attributed to the nutritional status. Overall, we found little effect of serial bleeding on blood cell production.

**Figure 1.**
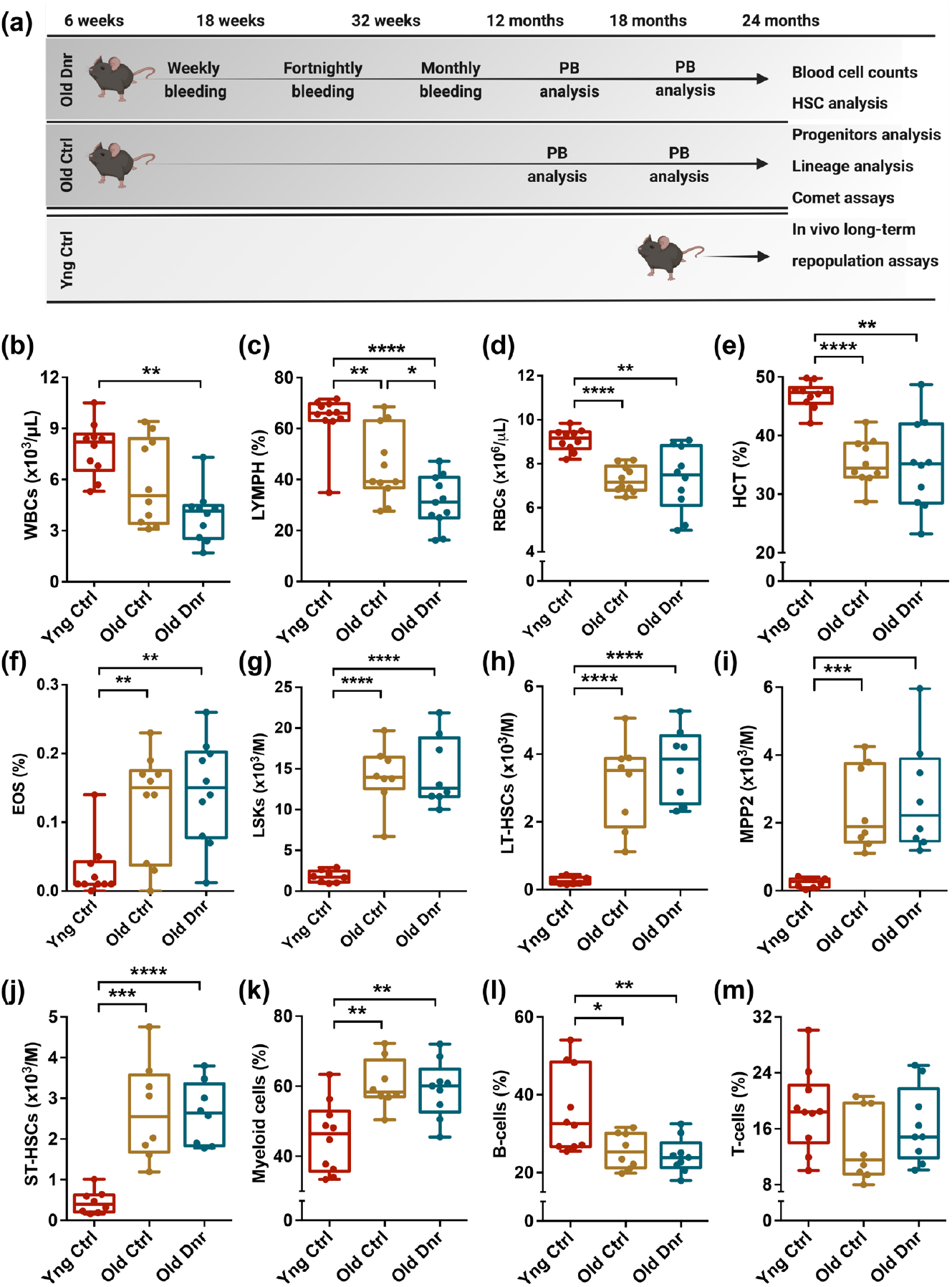
Serial bleeding does not affect the composition of hematopoietic population over aging. (a) Schematics depicting the experimental design. Six-weeks old mice were subjected to a long-term serial bleeding regimen. Detailed analysis of the hematopoietic system was followed to assess the effect of enhanced proliferative events on age-associated functional loss in HSCs. (b-f) 24-months old control and donor groups of mice were compared with 6-weeks old control mice for (b) WBCs, (c) lymphocytes, (d) RBCs, (e) hematocrit, and (f) eosinophiles. (g-j) Flow cytometry based analysis of various HSPC populations in the BM. Old donor mice that underwent serial bleeding regimen were compared with young and old control mice for the frequency of (g) LSKs (Lin^−^Sca-1^+^c-kit^+^ cells) (h) LT-HSCs (Lin^−^Sca-1^+^c-kit^+^CD150^+^CD48^−^ cells), (i) MPP2 (Lin^−^Sca-1^+^c-kit^+^CD150^+^CD48^+^), and (j) ST-HSCs (Lin^−^ Sca-1^+^c-kit^+^CD150^−^CD48^−^). (k-m) The three groups of mice compared for the proportion of (k) CD11b/Gr-1^+^ myeloid cells, (l) B220^+^ B-cells, (m) CD4/CD8^+^ T-cells in the BM.

Along with altered blood cell production, the function and composition of hematopoietic stem and progenitor cell (HSPC) population changes with age (*1, 13*). A robust increase in the HSC population identified as Lin^−^Sca-1^+^c-kit^+^ (LSK) side population (*14*), CD34^−^LSK cells (*15*) as well as CD48^−^CD34^−^ EPCR^+^CD150^+^LSK cells has been reported (*16*) with no change in short-term (ST-) HSCs (CD34^+^Flk2^−^ LSK cells) (*10*). Therefore, we next analyzed if serial bleeding regimen had any effect on the age-associated functional changes in HSC population (Fig. S2a). As compared with the young mice, both groups of old mice showed robust increase in the frequency of LSK cells (Fig. 1g), LT-HSCs (Fig. 1h,S2b), MPP2 (Fig. 1i,S2c), ST-HSCs (Fig. 1j,S2d), and MPP3/4 (Fig. S2e,f) populations. Interestingly, no effect of serial bleeding was observed when old donor mice were compared with old controls. In aged hematopoietic system, myeloid bias linked with altered differentiation potential of HSCs (*17*), or changes in lineage-committed progenitor populations (*18*) has been extensively reported. We performed flow cytometry analysis of lineage committed cells in the BM and compared the proportion of cells from myeloid and lymphoid lineage in the three groups of mice (Fig. S2g). Our experiments also showed robust increase in the myeloid cell population in the BM of the aged mice groups (Fig. 1k). Concomitantly, we noted a significant decrease in B-cell population (Fig. 1l), while no change was observed in the T-cell population (Fig. 1m). Hence, these experiments also showed no effect of serial bleeding regimen on the change in lineage composition of the hematopoietic system within the BM over aging.

In vivo data supported by long-term engraftment studies has also established a myeloid biased, poor hematopoietic reconstitution capacity of aged HSCs (*16, 19*). We used whole BM MNCs in our repopulation assays to assess if repeated hematopoietic insult induced by serial bleeding had any adverse impact on HSC aging (Fig. 2a). Up to a period of 8-weeks, we did not observe any decline in donor-derived engraftment resulting from aged HSPCs (Fig. 2b,c). However, from 12-week onwards post-transplantation, we noted a significant decrease in the donor-derived engraftment in the two aged mice groups (Fig. 2d,e), without any effect of serial bleeding regimen. We also noted a myeloid biased long-term engraftment following transplantation of aged HSPCs (Fig. 2f-h). These results are consistent with earlier studies that attributed this phenotype to an altered clonal diversity (*20*) and enrichment of myeloid committed progenitors (*21*).

**Figure 2.**
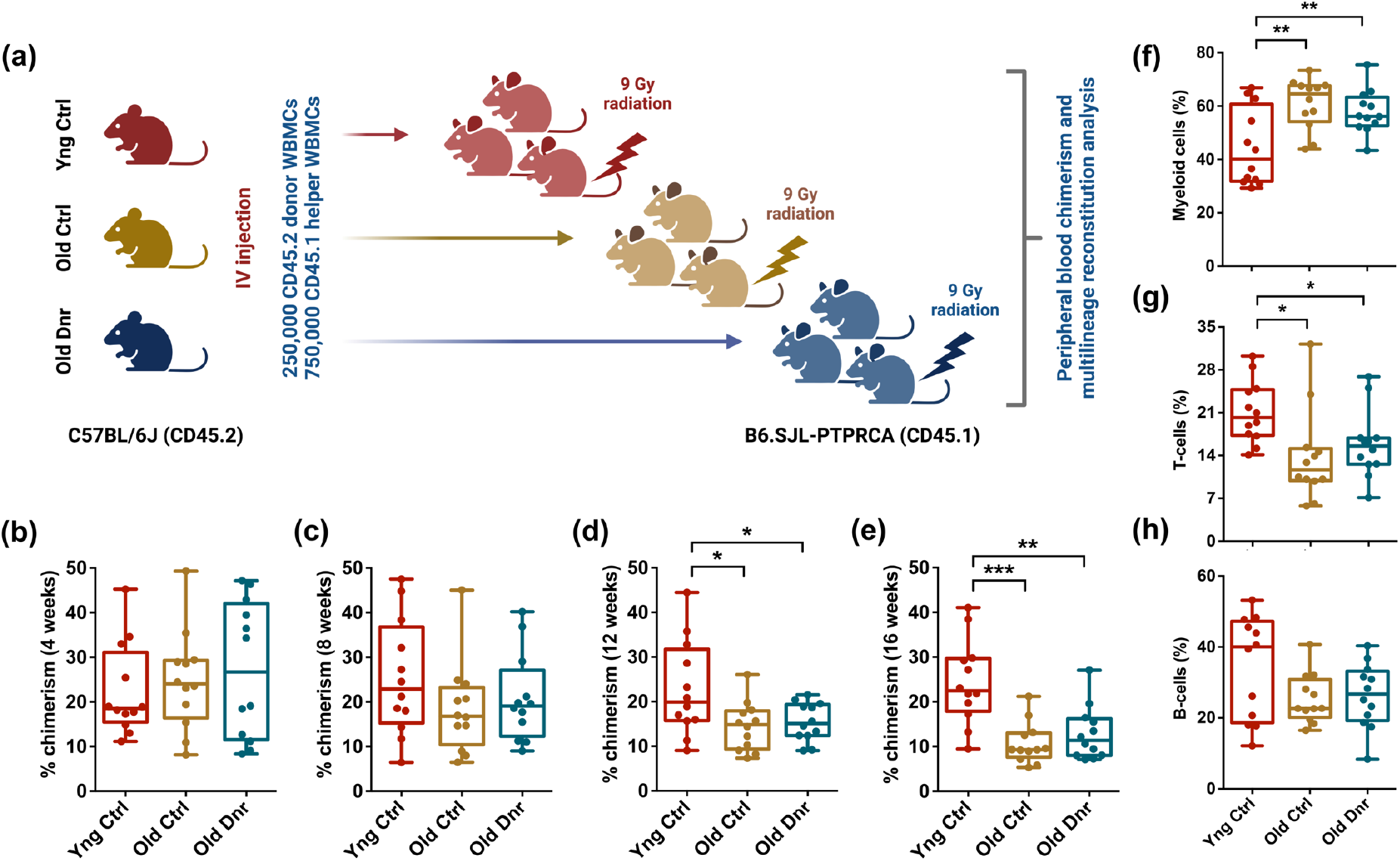
Repeated proliferative events in HSPCs during the lifetime do not alter long-term repopulation potential. (a) Schematic representation of the long-term hematopoietic reconstitution assays performed to compare the function of HSCs from old donor mice with young and old control mice. Donor derived (CD45.2) 250,000 whole BM cells were transplanted along with 750,000 (CD45.1) whole BM supporting cells into lethally irradiated (CD45.1) mice. Donor-derived peripheral blood chimerism was compared among the three groups of mice after (b) 4 weeks, (c) 8 weeks, (d) 12 weeks, and (e) 16 weeks of transplantation. (f-h) Comparison of the donor derived lineage engraftment in the three groups of mice. Flow cytometry analysis was performed to examine the contribution of donor derived cells within the (f) CD11b/Gr-1^+^ myeloid cell, (g) CD4/CD8^+^ T-cell, and (h) B220^+^ B-cell populations. (n=8-10), unpaired two-tailed Student’s t-test was performed. *p < 0.05, **p < 0.01, ***p < 0.001 and ****p < 0.0001.

Previous studies that reported DNA damage accumulation in aged HSCs (*22*), used comet assays performed under alkaline conditions for higher sensitivity (*23*). To examine the effect of extensive HSPC proliferation on the accumulation of DSBs specifically, we chose to perform neutral comet assays (*24*). As expected, the proportion of LSK cells with comet tail representing DSBs was significantly higher in the aged mice groups and remained unaffected by serial bleeding regimen (Fig. 3a,b). However, comparison of DSBs per cell between different groups in terms of Olive tail moment revealed a significant clearance of DNA damage following serial bleeding regimen (Fig. 3c). As proposed earlier, we attributed this clearance of DSBs to proliferative events during the lifetime (*7*). To further link DDR pathways with proliferation, we used available single-cell transcriptomic data from aged HSPCs (*25*). Principal Component Analysis (PCA) and Gene Set Enrichment Analysis (GSEA) were performed on proliferative and non-proliferative HSPCs identified using CellCycleScoring function of R-package Seurat (Fig. S3a). Analysis showed a robust increase in the overall enrichment of DDR genes (Fig. 3d, Table S1) and Hallmark pathways (Fig. S3b, Table S2) in proliferative compared to the quiescent HSPCs in aged mice. Interestingly, higher expression of DNA repair genes remained consistent for proliferative HSPCs from young mice as well (Fig. 3d). Reactome pathway analysis (Table S3) of upregulated DDR genes revealed a significant enrichment of DDR pathways in proliferative as compared with the quiescent HSPCs from aged mice (Fig. 3e). These results point towards the involvement of proliferation-coupled DDR pathways in clearing the DNA damage accumulated in HSPCs that undergo long dormancy period. We then used scRNA Seq data from young and aged human HSCs and compared the gene expression profiles of quiescent and proliferative HSCs (Fig. 3f). This analysis demonstrated extensive similarities between the mouse and human hematopoietic populations in terms of activation of DDR. We noted an increase in the enrichment of genes involved in DDR pathways (Fig. 3f, Table S4). This was also reflected in the Reactome pathways analysis and showed significant upregulation of DDR pathways in proliferative compared to the quiescent HSCs in humans (Fig. 3g, Table S5). Our previous results also have described enhanced DDR pathways and DNA repair in proliferative fetal liver HSPCs than adult BM HSCs (*26*). Next, we examined if proliferation induced by short-term serial bleeding regimen in the aged mice could have an impact on DDR gene expression and the status of accumulation of DNA damage. To this end, we first selected some of the DDR genes up-regulated in the proliferative HSPCs in young as well as aged mice (Fig. S3c, Table S1). Notably, most of these genes (*PCNA, RPA3, RFC4, RFC5, NEIL3, TOPBP1, BRCA1*) were also significantly (p<0.0001) up-regulated in the proliferative aged human HSCs (Fig. 3h, Table S4).

**Figure 3.**
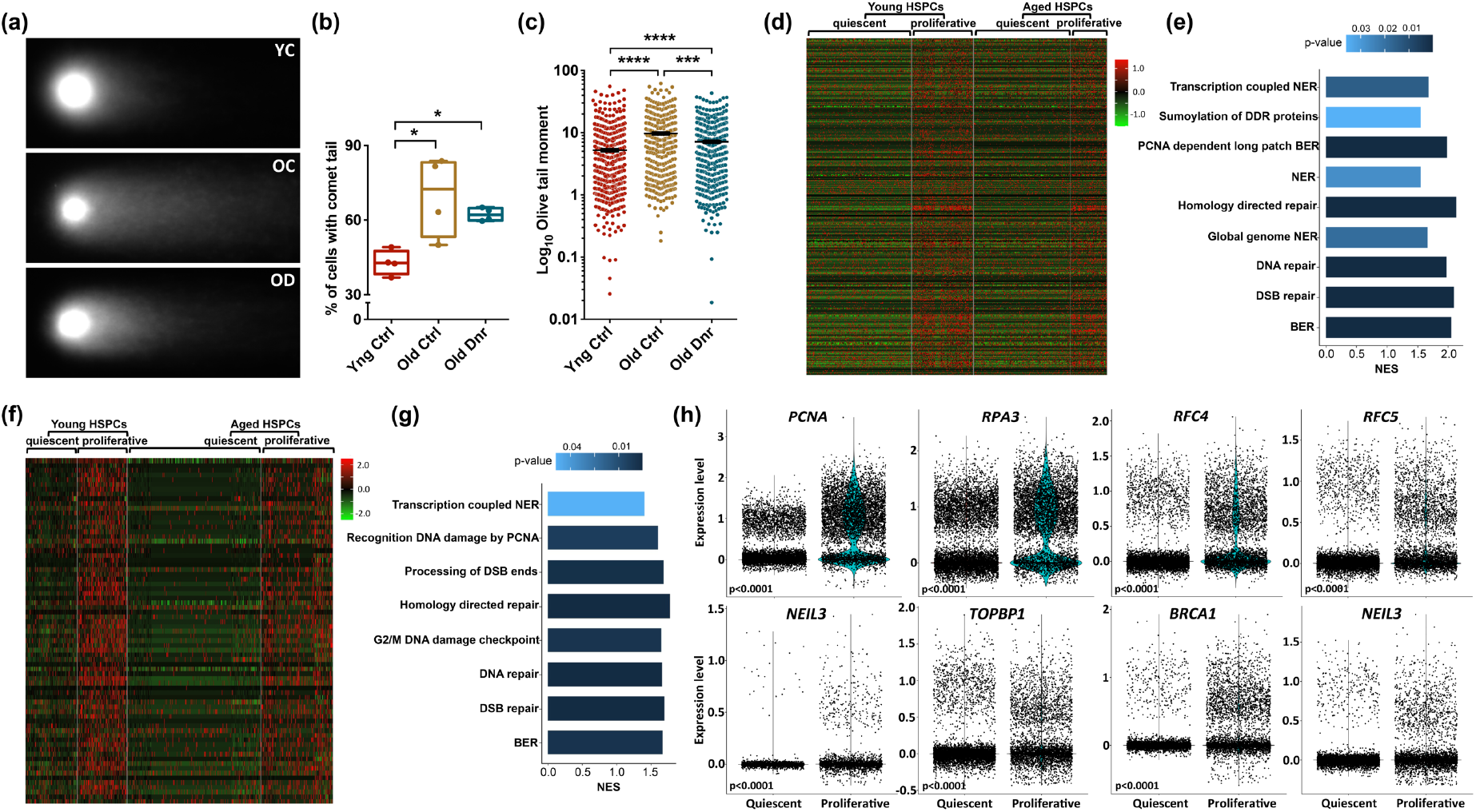
Proliferative events help in the repair of aging induced DNA damage. (a-c) Neutral comet assays performed to examine the level of DNA DSBs in HSPC population from old mice that underwent long-term serial bleeding regimen in comparison to young and old controls. (a) Representative comet images from freshly sorted LSK cells from the BM of mice from each group. (b) Proportion of LSK cells from the three groups of mice that showed comet tail. (c) Comparison of Olive tail moment as a measure of sheared DNA per cell. (d,e) Single cell RNA-Seq analysis to compare the gene expression and pathways enriched in quiescent versus proliferative mouse HSPCs using R-package Seurat. Data on FACS sorted Flt3^−^Lin^−^Sca-1^+^c-Kit^+^ cells from young and aged mice was acquired from public database (*25*). (d) Heatmap depicting expression of 409 genes associated with DNA damage response (DDR) in quiescent and proliferative HSPCs from young and old mice. (e) Differentially expressed Reactome DDR pathways identified using GSEA. The bars depict the normalized enrichment scores (NES) with p<0.05. NES>0 indicates signature enrichment in proliferative compared to quiescent HSPCs from aged mice. (f) Heatmap showing relative expression of 351 DDR pathway genes in quiescent and proliferative HSCs from young and aged humans. (g) Differentially enriched Reactome DDR pathways identified in proliferative compared to quiescent aged human HSCs using GSEA. The bars depict the normalized enrichment scores (NES) with p<0.05. NES>0 indicates signature enrichment in proliferative compared to quiescent HSCs from aged humans. (h) Violin plots depicting the feature gene expression of the selected genes associated with DNA damage repair pathways in quiescent and proliferative aged human HSCs.

We performed a short-term serial bleeding regimen in aged mice wherein the mice were bled thrice within a period of 10 days. We harvested lineage-depleted BM cells to examine the effect this proliferative regimen on the expression of DDR pathway genes by performing quantitative RT-PCR (Fig. 4a-d). Results clearly showed elevated transcript levels of most of the genes across the DDR pathways tested (mismatch repair or MMR; Fig. 4a, base excision repair or BER; Fig. 4b, homology-directed repair or HR; Fig. 4c, and nucleotide excision repair or NER; Fig. 4d). Finally, we examined if the enhanced DDR pathways could alter the status of DNA damage accumulated in aged HSPCs. We FACS sorted LSK cells from the mice that underwent short-term bleeding regimen and age-matched controls and performed neutral comet assays (Fig. 4e,f). These experiments provided striking results as we noted a massive decrease in the Olive tail moment showing clearance of DSBs (Fig. 4e,f). Overall, these results are in contrast to the prevalent notion that higher proliferation rates in HSCs can exacerbate DNA damage accumulation. Evidence also shows that the level of DNA damage accumulation might not be directly linked with age-associated functional decline. In contrast, we show that proliferative events in HSCs can activate DNA repair mechanisms to reduce the load of DNA damage acquired with age. As mutation accumulation is linked to several hematologic pathologies, serial bleeding as a simple regimen to clear DNA damages can have clinical implications.

**Figure 4.**
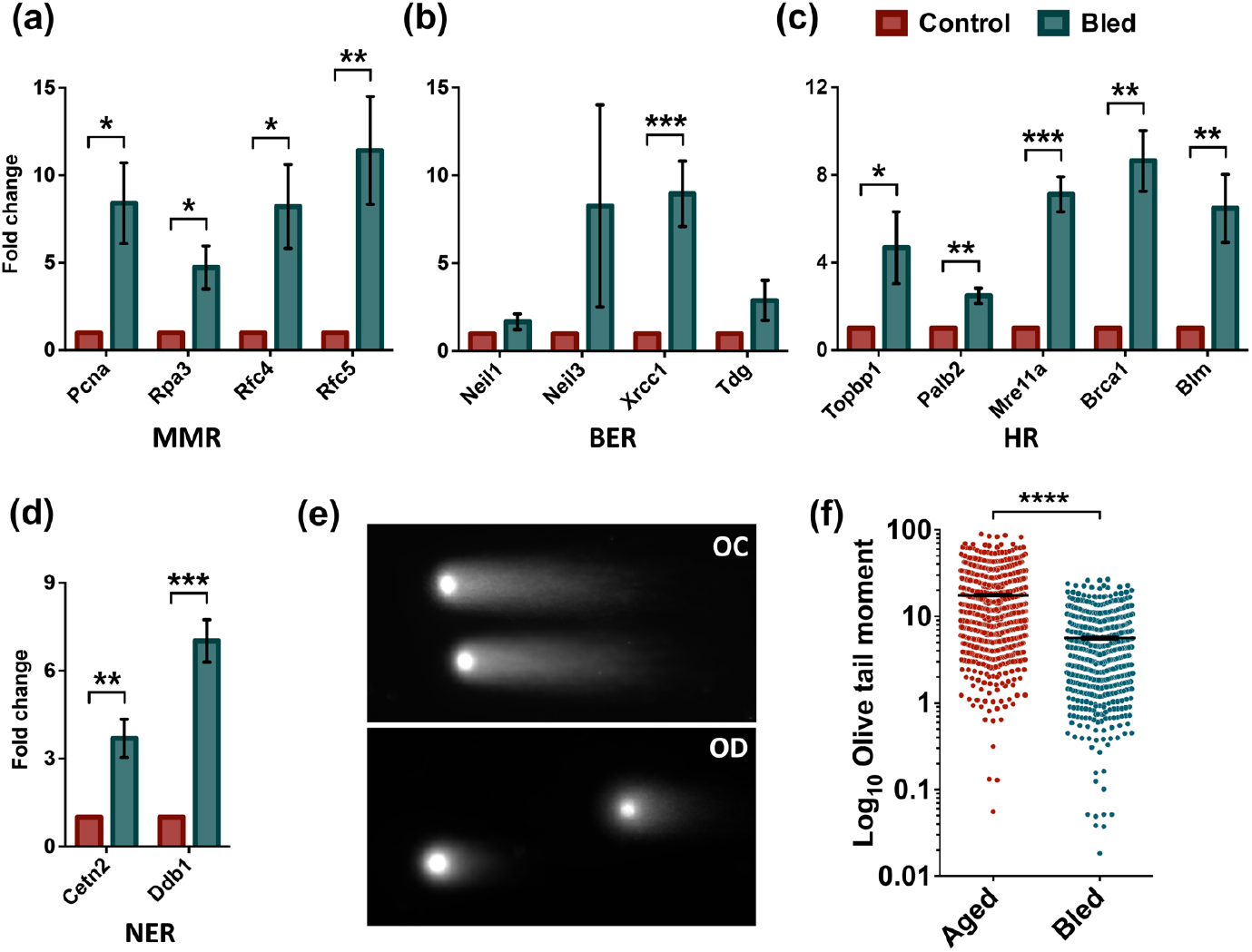
Short-term proliferative events help in the clearance of aging induced DNA damage. (a-d) Quantitative RT-PCR based analysis to examine the change in the expression of selected DDR pathway genes in BM derived lin-depleted cells following a short-term bleeding regimen in aged mice. The expression of genes with function in (a) mismatch repair or MMR (*Pcna, Rpa3, Rfc4* and *Rfc5*), (b) base excision repair or BER (*Neil1, Neil3, Xrcc1* and *Tdg*), (c) homologous recombination or HR (*Topbp1, Palb2, Mre11a, Brca1* and *Blm*), and (d) nucleotide excision repair or NER (*Cetn2, Ddb1*). (e,f) Neutral comet assay performed on FACS sorted LSK cells from aged mice with or without short-term bleeding regimen followed. (e) Representative comet images showing the level of DNA damage due to DSBs. (f) Comparison of olive tail moment from neutral comet assays performed on LSK cells from the two groups of mice. (n=3-5), unpaired two-tailed Student’s t-test was performed. *p < 0.05, **p < 0.01, ***p < 0.001 and ****p < 0.0001.

## Materials and Methods

### Mice

Six weeks to twenty-four months old C57BL/6J (CD45.2) and B6.SJL-PTPRCA (CD45.1) mice were bred and maintained in the animal facilities at KU Leuven, Belgium and IISER Thiruvananthapuram, India. During the experiments, mice were maintained in isolator cages at humidified constant temperature with a 12 h light-dark cycle. The mice were fed with autoclaved water and irradiated food (Safe diet, France) ad libitum. All animal experiments were approved by the Institutional Animal Ethics Committees for the respective animal facilities. At IISER Thiruvananthapuram, the animals were maintained as per guidelines provided by the Committee for the Purpose of Control and Supervision of Experiments on Animals (CPCSEA), Ministry of Environment and Forests, Government of India.

### Blood withdrawal

To induce excessive rounds of proliferation in HSCs during the lifetime, a long-term serial bleeding regimen was followed. Mice aged six weeks were used to draw 200µl blood via tail clipping method. Blood was drawn weekly for 12-weeks, followed by fortnightly bleeding for 14-weeks and monthly bleeding till 1 year of age. The mice were thereafter maintained for a period of two years and HSCs analysis was performed at 12 months, 18 months and 24 months of age. A short-term serial bleeding regimen of three blood withdrawals in 10-days period with three days intervals was used in aged mice (24 months of age) to induce HSC proliferation. At every time point, peripheral blood analysis was performed on Erba hematology analyzer for detailed blood cell counts. Detailed functional analysis of the hematopoietic system was performed at 24 months of age.

### Bone marrow aspiration

Mice were sacrificed via cervical dislocation and hindlimb bones were harvested. Adjacent muscle tissues were removed and bones were flushed with 1X PBS using a syringe with 26G needle. The resulting cell suspension was passed through a 41µm cell strainer (Corning, USA). The filtered cell suspension was diluted with 1X PBS and centrifuged at 600xg for 5 min at 4°C. The BM mononuclear cells were carefully resuspended in 1mL 1X PBS and were counted manually using a Neubauer hemocytometer (Neubauer, Germany).

### Transplantation

Frequency and function of the HSCs from young control, old control and old donor mice was examined using long-term hematopoietic reconstitution assays. Freshly isolated 250,000 whole BM cells (CD45.2^+^) were transplanted along with 750,000 competitor WBMCs (CD45.1^+^) into lethally irradiated (9Gy) 8-12 weeks old female mice. All irradiated mice were fed on enrofloxacin (Baytril) containing water. Peripheral blood chimerism and multilineage engraftment analysis was performed every 4 weeks by flow cytometry.

### Flow cytometry

Analysis of donor-derived chimerism, multi-lineage engraftment and characterization of hematopoietic system was performed by flow cytometry. For chimerism and multi-lineage engraftment, donor and recipient cells were identified as CD45.2^+^ and CD45.1^+^ cells, respectively. Within the CD45.2^+^ donor derived cells, lineage-committed cells were identified as myeloid (CD11b^+^/Gr-1^+^), T-cell (CD3e^+^) and B-cell (B220^+^) populations. On the basis of the expression of SLAM markers, CD150 and CD48, lin^−^c-kit^+^Sca-1^+^ (LSK) population was subdivided into four subpopulations: CD150^+^CD48^−^ (LT-HSCs), CD150^+^CD48^+^ (MPP2), CD150^−^CD48^+^ (MPP3/4), and CD150^−^CD48^−^ (ST-HSCs) were characterized by allophycocyanin-conjugated anti-lineage antibody cocktail, PE-conjugated anti-mouse c-kit antibodies, BB700-conjugated anti-mouse Sca-1 antibodies, FITC-conjugated anti-mouse CD48 antibodies, and PECy7-conjugated CD150 antibodies. Suitable isotype controls for each antibody were used in all experiments. A complete list of antibodies used for these experiments is provided in the Table S6.

Cell cycle analysis was performed on BM derived MNC population to examine the effect of serial bleeding regimen on HSPC proliferation. The BM MNCs were first immuno-labelled for identification of LSK cells, as described above. After cell surface staining with FITC conjugated anti-lineage antibody cocktail, BB700 conjugated anti-mouse Sca-1 and PE conjugated anti-mouse c-kit antibodies, the cells were fixed using BD Cytofix/Cytoperm buffer. The cells were then washed with Perm/Wash buffer and incubated with AF647 conjugated Ki-67 antibodies. The cells were further washed and labelled with DAPI for 30 mins on ice. The samples were acquired using FACS Aria III (BD Biosciences, San Jose, CA) and FlowJo software (TreeStar, Ashland, OR).

### Neutral comet assay

This method was used to perform comet assay to assess DNA damage in freshly sorted BM derived HSPCs under neutral pH. FACS sorted LSK cells were resuspended in low-melting agarose (Sigma; type VII). The suspension was poured onto agarose-coated comet slides dropwise and incubated at 4°C. After agarose solidifies, slides were immersed in lysis buffer (2.5 M NaCl, 0.1 M EDTA, 10 mM Trizma base, 1% Triton X-100, 10% DMSO, pH 10) for 1h. The slides are then washed with pre-chilled neutral electrophoresis buffer and incubated for 30 minutes. The slides are then transferred for electrophoresis in pre-chilled 1X neutral electrophoresis buffer (10X buffer contains 1M Trizma base, 3M Sodium acetate trihydrate, pH is set at 9 using glacial acetic acid). After electrophoresis (∼60 min, 21 V), air-dried and precipitated slides (10X precipitation buffer contains 7.5M ammonium acetate, which is diluted to 1M using 100% ethanol) were stained with SYBR-Gold solution (1:1000). Olive Tail Moment was scored for 100-200 cells/sample by using Open Comet plugin in ImageJ software.

### Quantitative RT-PCR

Hematopoietic progenitors were isolated with EasySep™ mouse hematopoietic progenitor isolation kit. Total RNA was prepared using TRIzol™ Reagent. The purity and the concentration of RNA were assessed using a micro-volume spectrophotometer (Colibri, Berthold Technologies GmBH & Co. KG Germany). Two micrograms of RNA from each sample was used to synthesize cDNA using PrimeScript RT reagent kit (Takara Biotechnology Co. Ltd. Dalin, China) according to the manufacturer’s protocol. Quantitative PCR was carried out using TB Green Premix (Takara Biotechnology Co. Ltd. Dalin, China). The PCR reactions were carried out in a CFX96 detection system (Thermal-Cycler C1000, Bio-Rad Laboratories, Hercules, CA, USA). The list of primers used is given in Table S7.

### Single-cell RNA Seq analysis

Single cell RNA seq libraries of BM HSPCs isolated from young and aged mice were retrieved from the Gene Expression Omnibus (GEO; GSE147729) (*25*) and human HSCs scRNA libraries from (GSE180298) (*27*). The gene count expression matrix was analyzed using the Seurat v.4.1.1 in RStudio. Cells containing fewer than 100 genes and fewer than 500 unique molecular identifiers were excluded from subsequent analysis. Normalization of raw counts was performed using the NormalizeData function in Seurat. Variable genes were identified using the FindVariableGenes function. Expression values in the dataset were scaled and centered for dimensional reduction using the ScaleData function in Seurat, with default parameters. Principal Component Analysis (PCA) was performed following analysis of cell cycle phase for individual cell was performed with CellCycleScoring function of Seurat, employing a core gene set as previously described (*28*), distinguishing between cells in different stages. As HSCs at G_o_ stage are transcriptionally closer to G_1_ than S or G_2_/M cells we marked them quiescent, while HSCs in the S and G_2_/M phases were identified as proliferative. Differential gene expression analysis was performed with FindMarkers function. Subsequently, Hallmark and Reactome pathway enrichment analysis was performed with the differential gene expression dataset, employing a fast gene set enrichment analysis package. Pathways demonstrating statistical significance, indicated by a p-value ≤ 0.05, were considered noteworthy.

### Quantification and statistical analysis

Data are represented mean±SEM or as box and whisker plots (Min to Max). Comparisons between samples from two groups with normally distributed data with equal variance were made using the unpaired two-tailed Student’s t-test. Mann Whitney test was used for comparing two groups where data were non-normally distributed. Statistical analyses were performed with Microsoft Excel, GraphPad Prism 6 and RStudio. For all analyses, p-values <0.05 were accepted as statistically significant.

## Author Contributions

SK conceptualized, designed and supervised the study, wrote the manuscript. SHM and IMR performed the experiments, analyzed the results and reviewed the manuscript. HA performed computational analysis on mouse and human single cell RNA-Seq data, reviewed the manuscript. SS provided technical support for experiments on animals.

## Acknowledgements

This work was supported by the DBT/Wellcome Trust India Alliance Fellowship (IA/I/15/2/502061) awarded to SK and intramural funds from Indian Institute of Science Education and Research Thiruvananthapuram (IISER TVM). IISER TVM Institutional animal facility is supported by funds from the Department of Science and Technology, Government of India (under FIST scheme; SR/FST/LS-II/2018/217). The authors wish to thank Prof. Catherine Verfaillie, KU Leuven for providing material support and hosting SK for performing experiments on animals. SHM and HA are supported by IISER TVM. IMR was supported by the senior research fellowship from University Grants Commission (UGC), India.

## Data availability

The published datasets used for the analysis were retrieved from the GEO using the accession number GSE147729 (*25*) and GSE180298 (*27*).

## Code availability

The code for the bioinformatics analysis, alongside information on software and package versions, is available at https://github.com/stemcellbiologylab.

---

**Figure S1.**
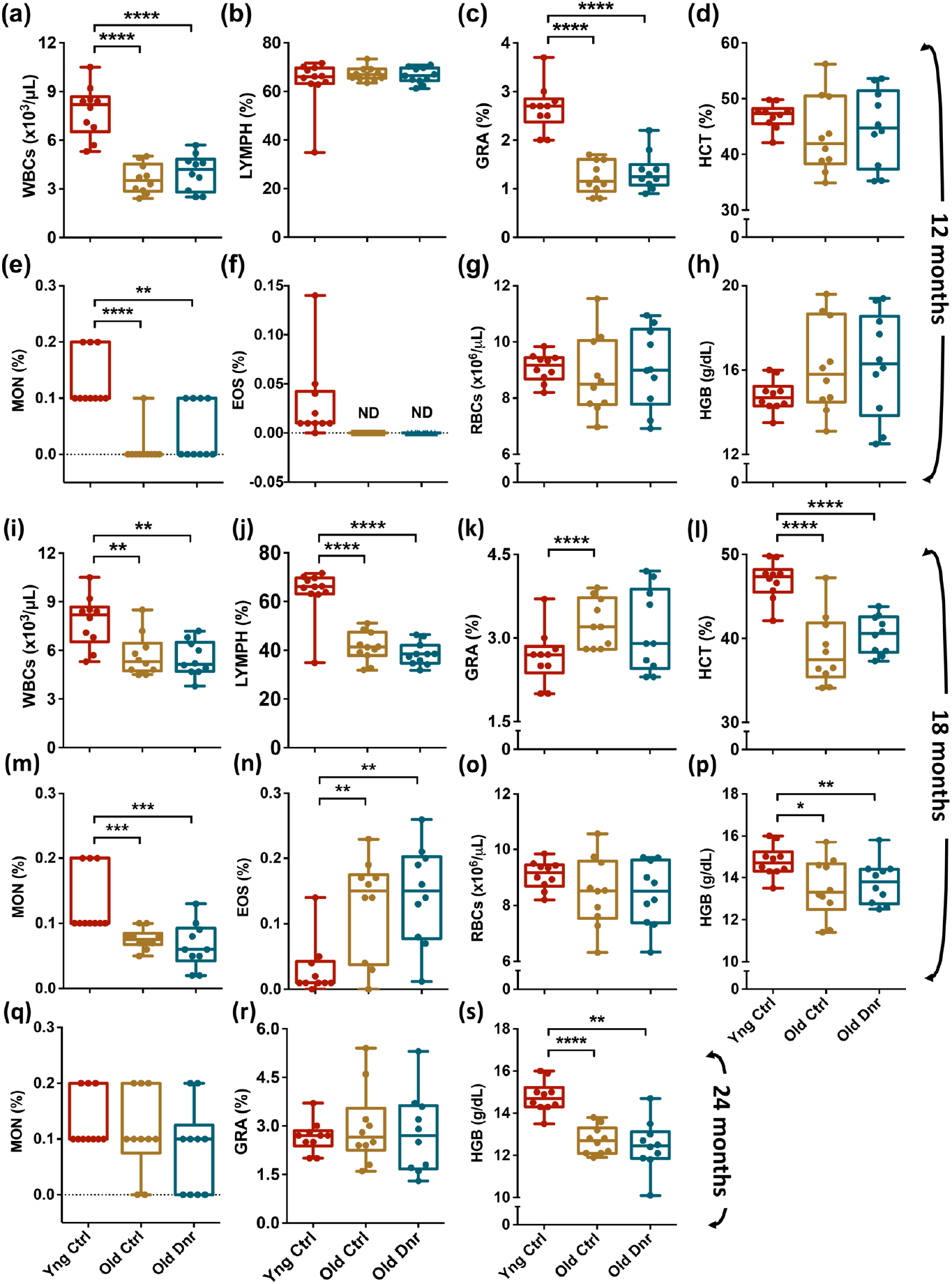
Serial bleeding induces HSC proliferation without affecting blood cell production. (a-h) Peripheral blood cell counts from the two groups of 12-months old mice compared with young adults. The peripheral blood from the mice was analysed for (a) WBCs, (b) lymphocytes, (c) granulocytes, (d) hematocrit, (e) monocytes, (f) eosinophils, (g) RBCs, and (h) hemoglobin. (i-p) Peripheral blood cell counts from 18 months old control and donor groups in comparison to the 6-weeks old mice. The peripheral blood was used to compare (i) WBCs, (j) lymphocytes, (k) granulocytes, (l) hematocrit, (m) monocytes, (n) eosinophils, (o) RBCs, and (p) hemoglobin. (q-s) Peripheral blood cell counts from 24 months aged old control and donor groups were analyzed along with six-weeks old mice for (q) monocytes, (r) granulocytes, and (s) hemoglobin. (n=5-10), Unpaired two-tailed Student’s t-test was performed. *p < 0.05, **p < 0.01, ***p < 0.001 and **** < 0.0001.

**Figure S2.**
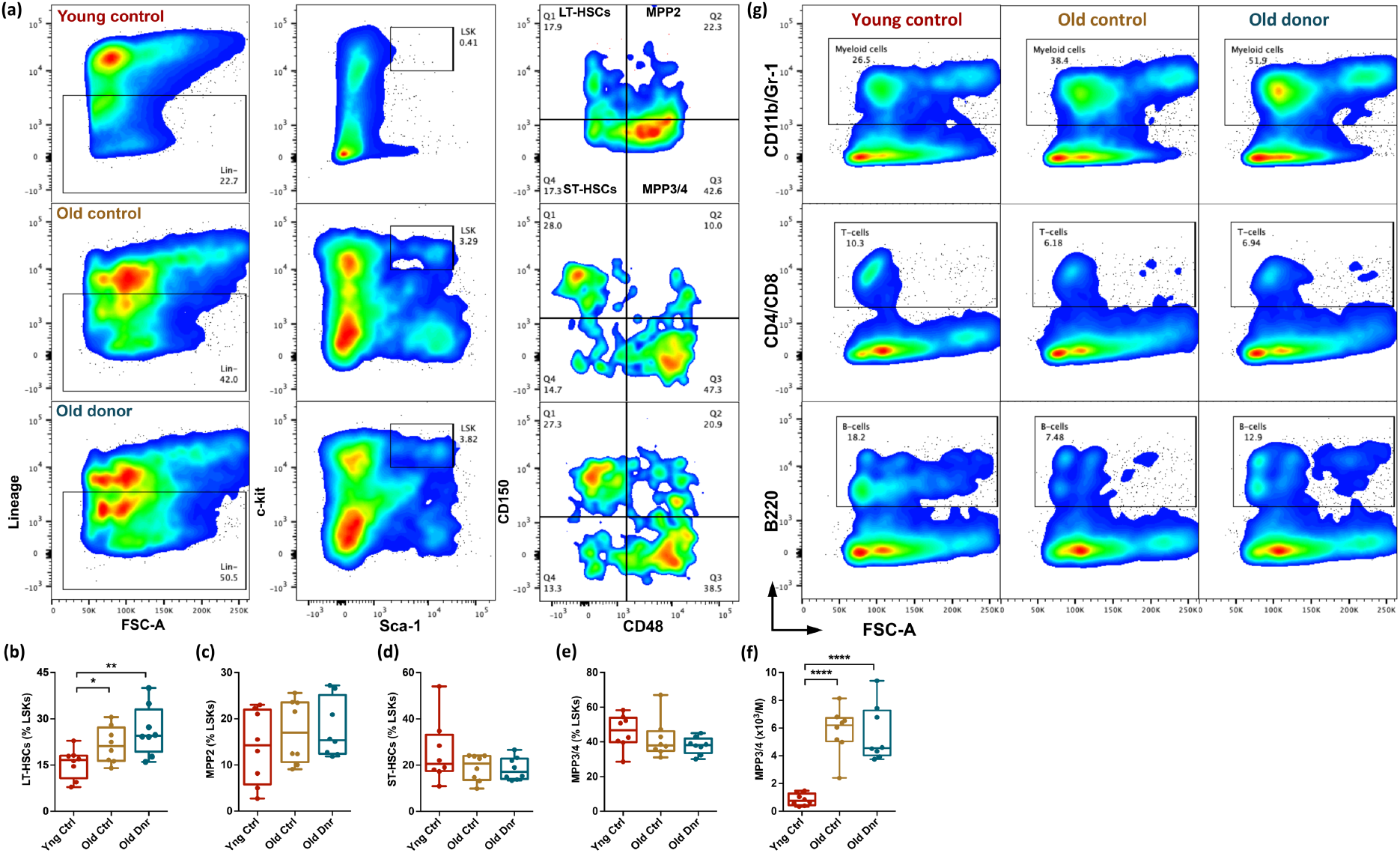
Serial bleeding does not affect the composition of hematopoietic stem and progenitor cell populations. (a-f) Bone marrow derived MNCs from 24 months old control and donor mice was compared with young adult (6-weeks old) mice for hematopoietic stem and progenitor populations (HSPCs). The three groups of mice were analyzed for the frequency of HSPCs in the BM MNC population. (a) Flow cytometry profiles to analyze LT-HSC (Lin^−^Sca-1^+^c-kit^+^CD150^+^CD48^−^), ST-HSCs (Lin^−^Sca-1^+^c-kit^+^CD150^−^ CD48^−^), MPP2 (Lin^−^Sca-1^+^c-kit^+^CD150^+^CD48^+^) and MPP3/4 (Lin^−^Sca-1^+^c-kit^+^CD150^−^CD48^+^) populations. (b-e) The LSK population in the BM MNCs further analyzed for the frequency of (b) LT-HSCs, (c) MPP2, (d) ST-HSCs, and (e) MPP3/4. The proportion of these cell populations within the LSK population has been shown. (f) The frequency of MPP3/4 (per million) within the total BM MNC population. (g) The bone marrow derived MNCs from old control and donor mice groups along with young mice analyzed for the proportion of lineage committed cells. Flow cytometry analysis performed on the BM MNC population to quantify myeloid (CD11b/Gr-1^+^; upper panel), T-cells (CD4/CD8^+^; middle panel), and B-cells (B220^+^; lower panel). n=8-10, Unpaired two-tailed Student’s t-test was performed. *p < 0.05, **p < 0.01, ***p < 0.001 and ****p < 0.0001.

**Figure S3.**
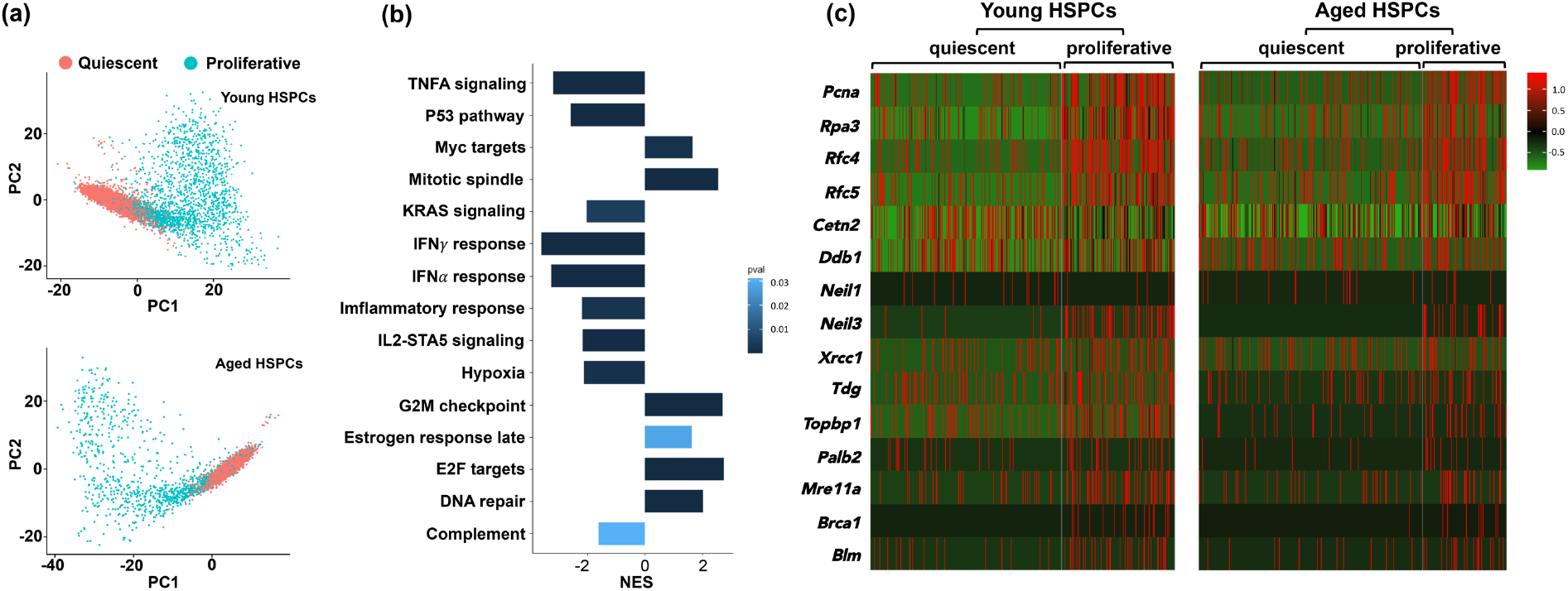
Induction of proliferation in HSCs enhances DDR. (a) Principal Component Analysis (PCA) based on first two components with highest variance. The plots show clustering of cells based on the expression of proliferation markers in HSPCs harvested from young (upper panel) and aged (lower panel) mice BM. (b) Hallmark Pathway analysis in proliferative compared to non-proliferative HSPCs based on single-cell RNA Seq data. Gene set enrichment analysis for MSigDB Hallmark 2023.2 pathways associated with aged HSPCs was performed. The bar depicts the normalized enrichment scores (NES) with p<0.05. NES>0 indicates signature enrichment in proliferative aged HSPCs compared to quiescent aged HSPCs, while NES<0 indicates enrichment in quiescent HSCs. (c) Heatmap depicting the feature gene expression of the selected 15 genes associated with DNA damage repair pathways in quiescent and proliferative HSPCs from young and aged mice. The selected genes belonged to different DDR pathways; mismatch repair (*Pcna, Rpa3, Rfc4* and *Rfc5*), base excision repair (*Neil1, Neil3, Xrcc1* and *Tdg*), homologous recombination (*Topbp1, Palb2, Mre11a, Brca1* and *Blm*) and nucleotide excision repair (*Cetn2* and *Ddb1*).

## References

1. S. J. Morrison, A. M. Wandycz, K. Akashi, A. Globerson, I. L. Weissman, The aging of hematopoietic stem cells. Nature medicine 2, 1011–1016 (1996).

2. S. Khurana, The Effects of Proliferation and DNA Damage on Hematopoietic Stem Cell Function Determine Aging. Dev Dynam 245, 739–750 (2016).

3. D. J. Rossi, D. Bryder, J. Seita, A. Nussenzweig, J. Hoeijmakers, I. L. Weissman, Deficiencies in DNA damage repair limit the function of haematopoietic stem cells with age. Nature 447, 725–729 (2007).

4. K. Ito, A. Hirao, F. Arai, S. Matsuoka, K. Takubo, I. Hamaguchi, K. Nomiyama, K. Hosokawa, K. Sakurada, N. Nakagata, Y. Ikeda, T. W. Mak, T. Suda, Regulation of oxidative stress by ATM is required for self-renewal of haematopoietic stem cells. Nature 431, 997–1002 (2004).

5. D. Walter, A. Lier, A. Geiselhart, F. B. Thalheimer, S. Huntscha, M. C. Sobotta, B. Moehrle, D. Brocks, I. Bayindir, P. Kaschutnig, K. Muedder, C. Klein, A. Jauch, T. Schroeder, H. Geiger, T. P. Dick, T. Holland-Letz, P. Schmezer, S. W. Lane, M. A. Rieger, M. A. Essers, D. A. Williams, A. Trumpp, M. D. Milsom, Exit from dormancy provokes DNA-damage-induced attrition in haematopoietic stem cells. Nature 520, 549–552 (2015).

6. J. Flach, S. T. Bakker, M. Mohrin, P. C. Conroy, E. M. Pietras, D. Reynaud, S. Alvarez, M. E. Diolaiti, F. Ugarte, E. C. Forsberg, M. M. Le Beau, B. A. Stohr, J. Mendez, C. G. Morrison, E. Passegue, Replication stress is a potent driver of functional decline in ageing haematopoietic stem cells. Nature 512, 198–202 (2014).

7. I. Beerman, J. Seita, M. A. Inlay, I. L. Weissman, D. J. Rossi, Quiescent hematopoietic stem cells accumulate DNA damage during aging that is repaired upon entry into cell cycle. Cell Stem Cell 15, 37–50 (2014).

8. S. H. Cheshier, S. S. Prohaska, I. L. Weissman, The effect of bleeding on hematopoietic stem cell cycling and self-renewal. Stem Cells Dev 16, 707–717 (2007).

9. L. Ferrucci, A. Corsi, F. Lauretani, S. Bandinelli, B. Bartali, D. D. Taub, J. M. Guralnik, D. L. Longo, The origins of age-related proinflammatory state. Blood 105, 2294–2299 (2005).

10. M. C. Florian, K. Dorr, A. Niebel, D. Daria, H. Schrezenmeier, M. Rojewski, M. D. Filippi, A. Hasenberg, M. Gunzer, K. Scharffetter-Kochanek, Y. Zheng, H. Geiger, Cdc42 activity regulates hematopoietic stem cell aging and rejuvenation. Cell Stem Cell 10, 520–530 (2012).

11. A. Rundberg Nilsson, S. Soneji, S. Adolfsson, D. Bryder, C. J. Pronk, Human and Murine Hematopoietic Stem Cell Aging Is Associated with Functional Impairments and Intrinsic Megakaryocytic/Erythroid Bias. PloS one 11, e0158369 (2016).

12. M. Tettamanti, U. Lucca, F. Gandini, A. Recchia, P. Mosconi, G. Apolone, A. Nobili, M. V. Tallone, P. Detoma, A. Giacomin, M. Clerico, P. Tempia, L. Savoia, G. Fasolo, L. Ponchio, M. G. Della Porta, E. Riva, Prevalence, incidence and types of mild anemia in the elderly: the “Health and Anemia” population-based study. Haematologica 95, 1849–1856 (2010).

13. H. Geiger, G. de Haan, M. C. Florian, The ageing haematopoietic stem cell compartment. Nat Rev Immunol 13, 376–389 (2013).

14. S. M. Chambers, C. A. Shaw, C. Gatza, C. J. Fisk, L. A. Donehower, M. A. Goodell, Aging hematopoietic stem cells decline in function and exhibit epigenetic dysregulation. PLoS Biol 5, e201 (2007).

15. S. Noda, H. Ichikawa, H. Miyoshi, Hematopoietic stem cell aging is associated with functional decline and delayed cell cycle progression. Biochemical and biophysical research communications 383, 210–215 (2009).

16. B. Dykstra, S. Olthof, J. Schreuder, M. Ritsema, G. de Haan, Clonal analysis reveals multiple functional defects of aged murine hematopoietic stem cells. The Journal of experimental medicine 208, 2691–2703 (2011).

17. D. J. Rossi, D. Bryder, J. M. Zahn, H. Ahlenius, R. Sonu, A. J. Wagers, I. L. Weissman, Cell intrinsic alterations underlie hematopoietic stem cell aging. Proc Natl Acad Sci U S A 102, 9194–9199 (2005).

18. W. W. Pang, E. A. Price, D. Sahoo, I. Beerman, W. J. Maloney, D. J. Rossi, S. L. Schrier, I. L. Weissman, Human bone marrow hematopoietic stem cells are increased in frequency and myeloid-biased with age. Proc Natl Acad Sci U S A 108, 20012–20017 (2011).

19. T. T. Ho, P. V. Dellorusso, E. V. Verovskaya, S. T. Bakker, J. Flach, L. K. Smith, P. B. Ventura, O. M. Lansinger, A. Herault, S. Y. Zhang, Y. A. Kang, C. A. Mitchell, S. A. Villeda, E. Passegue, Aged hematopoietic stem cells are refractory to bloodborne systemic rejuvenation interventions. The Journal of experimental medicine 218, (2021).

20. C. E. Muller-Sieburg, H. B. Sieburg, Clonal diversity of the stem cell compartment. Curr Opin Hematol 13, 243–248 (2006).

21. C. Gekas, T. Graf, CD41 expression marks myeloid-biased adult hematopoietic stem cells and increases with age. Blood 121, 4463–4472 (2013).

22. B. M. Moehrle, K. Nattamai, A. Brown, M. C. Florian, M. Ryan, M. Vogel, C. Bliederhaeuser, K. Soller, D. R. Prows, A. Abdollahi, D. Schleimer, D. Walter, M. D. Milsom, P. Stambrook, M. Porteus, H. Geiger, Stem Cell-Specific Mechanisms Ensure Genomic Fidelity within HSCs and upon Aging of HSCs. Cell Rep 13, 2412–2424 (2015).

23. P. L. Olive, J. P. Banath, The comet assay: a method to measure DNA damage in individual cells. Nat Protoc 1, 23–29 (2006).

24. P. L. Olive, D. Wlodek, J. P. Banath, DNA double-strand breaks measured in individual cells subjected to gel electrophoresis. Cancer Res 51, 4671–4676 (1991).

25. L. Hérault, M. Poplineau, A. Mazuel, N. Platet, É. Remy, E. Duprez, Single-cell RNA-seq reveals a concomitant delay in differentiation and cell cycle of aged hematopoietic stem cells. BMC Biology 19, 19 (2021).

26. A. Biswas, I. M. Roy, P. C. Babu, J. Manesia, S. Schouteden, V. Vijayakurup, R. J. Anto, J. Huelsken, A. Lacy-Hulbert, C. M. Verfaillie, S. Khurana, The Periostin/Integrin-αv Axis Regulates the Size of Hematopoietic Stem Cell Pool in the Fetal Liver. Stem Cell Reports 15, 340–357 (2020).

27. M. Ainciburu, T. Ezponda, N. Berastegui, A. Alfonso-Pierola, A. Vilas-Zornoza, P. San Martin-Uriz, D. Alignani, J. Lamo-Espinosa, M. San-Julian, T. Jiménez-Solas, F. Lopez, S. Muntion, F. Sanchez-Guijo, A. Molero, J. Montoro, G. Serrano, A. Diaz-Mazkiaran, M. Lasaga, D. Gomez-Cabrero, M. Diez-Campelo, D. Valcarcel, M. Hernaez, J. P. Romero, F. Prosper, Uncovering perturbations in human hematopoiesis associated with healthy aging and myeloid malignancies at single-cell resolution. eLife 12, e79363 (2023).

28. X. Fan, J. Dong, S. Zhong, Y. Wei, Q. Wu, L. Yan, J. Yong, L. Sun, X. Wang, Y. Zhao, W. Wang, J. Yan, X. Wang, J. Qiao, F. Tang, Spatial transcriptomic survey of human embryonic cerebral cortex by single-cell RNA-seq analysis. Cell Res 28, 730–745 (2018).

